# Modulation of neurofluid fluctuation frequency by baseline carbon dioxide in awake humans: the role of the autonomic nervous system

**DOI:** 10.1101/2024.06.05.597472

**Authors:** Xiaole Z. Zhong, Catie Chang, J. Jean Chen

## Abstract

An understanding of neurofluid dynamics has been gaining importance, in part given the link between neurofluid dynamics and glymphatic flow. Recently, CSF pulsations have been attributed to widespread changes in cerebral blood volume (CBV) driven by sleep-state slow-wave electrocortical activity (Fultz et al., 2019), by localized neuronal activity (Williams et al., 2023), by respiration-related autonomic tone (Picchioni et al., 2022) and by vigilance (Z. Yang et al., 2024). It was further suggested that the slow-wave induced CSF pulsations are in fact driven by autonomic (ANS) regulation (Picchioni et al., 2022), and that CSF dynamics are ultimately modulated by ANS mechanisms instead of by sleep per se. To further understand the role of this ANS regulation of vascular tone independently of sleep, and given the established influence of carbon dioxide (CO_2_) on both ANS tone and vascular tone, we hypothesized that a modulation of basal CO_2_, producing altered global vascular tone and respiration, may highlight the role of ANS regulation in driving CSF flow, and more broadly, neurofluid flow. In this work, we report on observations of neurofluid dynamics at awake normocapnia as well as mild hyper- and hypocapnia steady states. We use the resting-state BOLD fMRI time courses in neurofluid regions (i.e. blood vessels, CSF compartments) as a surrogate of neurofluid dynamics. We found that 1) the manner biomechanical does not drive the variations in neurfluid dynamics across capnias; 2) besides respiration, cardiac pulsation also independently drives neurofluid flow as an indication of the ANS pathway of control; 3) changed CO_2_ alters neurofluid dynamics primarily through frequency rather than amplitude of heart-rate and respiratory-volume variability. These findings suggest that hyper- and hypocapnia both represent a disruption of homeostasis that engages ANS regulation, as reflected by the deviations in CRF and RRF from normocapnia. Our work demonstrates in awake humans previously reported ANS regulation observed during sleep. As basal CO_2_ can modulate this ANS regulation, it represents a new avenue for modulating neurofluid dynamics independently of sleep, attention or neuronal activation. More broadly, individuals with different basal capnic states may manifest differences in CSF dynamics, giving rise to a novel paradigm for modulating neurofluid flow in awake humans.

## Introduction

An understanding of neurofluid dynamics has been gaining importance, in part given the link between neurofluid dynamics and glymphatic flow (Taoka & Naganawa, 2020), which has in turn been linked to dementia pathology (Song et al., 2023). Neurofluids are defined as “fluids in which the central nervous system is immersed, such as blood, cerebrospinal fluid (CSF) and interstitial fluid (ISF)” (Taoka & Naganawa, 2020). Based on the current understanding, the drivers of CSF flow are arterial pulsation. These fluctuations can be further subdivided into vasomotion, respiration and cardiac activity (Rasmussen et al., 2022). The amplitude of CSF fluctuations have been related to glymphatic flow, but the exact interaction between CSF fluctuations and glymphatic flow remains unclear. It has been suggested that, in addition to CSF fluctuation amplitude, CSF fluctuation frequency is also of interest. The slow-wave CSF fluctuation that drives glymphatic flow is driven by vascular oscillators which can be further subdivided into endogenic (0.001-0.02 Hz), neurogenic (0.02-0.04 Hz) and vasogenic (0.06-0.14 Hz) (Bracic & Stefanovska, 1998). Myogenic oscillations are driven by Mayer waves, while neurogenic oscillations are primarily the result of innervation of the autonomic nervous system on the microvasculature (Strik et al., 2002).

Being sensitive to blood and CSF flow, blood-oxygenation-level-dependent (BOLD) resting-state fMRI (rs-fMRI) has recently enhanced the feasibility of broadening the study neurofluid dynamics (Diorio et al., 2023; Fultz et al., 2019; H.-C. S. Yang et al., 2022), although BOLD rs-fMRI’s primary application is mapping brain functional connectivity (FC) in health and disease. Using rs-fMRI, recent studies have highlighted the strong modulatory effects of sleep on CSF flow (Fultz et al., 2019), and in the process cementing the utility of blood-oxygenation-level-dependent (BOLD) resting-state fMRI (rs-fMRI) for monitoring CSF dynamics. Specifically, Fultz et al. attributed CSF pulsations to widespread changes in cerebral blood volume (CBV) driven by sleep-state slow-wave electrocortical activity in predominantly N1 and N2 sleep stages (Fultz et al., 2019). Indeed, CBV and CSF oscillations, two main types of neurofluid dynamics, were recently found to be linked through a temporal derivative (H.-C. S. Yang et al., 2022). Interestingly, Williams et al. later found CSF flow to be coupled to localized visual activity in the awake state, cementing the role of electrocortical activity irrespective of sleep (Williams et al., 2023). However, widespread CBV (and by inference CSF flow) modulations can be achieved in the awake state through myoactive mechanisms independently of electrocortical activity, including by modulating intravascular carbon dioxide (CO_2_).

The primary biomechanical driver of CBV fluctuations and hence CSF flow remains arterial pulsation (Rasmussen et al., 2022), it is reasonable to expect CSF fluctuation amplitude to be related to vascular tone. That is, the manner in which the rhythmic pressure waves generated by the heartbeat in the arteries are transmitted and modified in the CSF system depends on biomechanical factors such as brain-tissue compliance and vascular reactivity. Thus, at reduced vascular tone, the ability of CBV to respond to arterial pulsation is reduced, resulting in reduced ability to induce CSF fluctuations. CO_2_ is a potent vasodilator, and CO_2_ modulations have been used to achieve vascular tone modulation (Xie et al., 2006). Hypercapnia can achieve vasodilation (Kety & Schmidt, 1948; Reivich, 1964), and hypocapnia vasoconstriction. Both hyper- and hypocapnia result in diminished vascular tone and hence reduced BOLD signal amplitude (Halani et al., 2015). This reduced vascular tone can also be reflected in changes in the frequency characteristics of the rs-fMRI response (E. R. Cohen et al., 2002). Moreover, hypercapnia is known to increase CBV and decrease CSF volume (van der Kleij et al., 2020). Thus, there is ample evidence to support hypercapnia and hypocapnia modulating brain hemodynamics. This is analogous to the case of hypertension, whereby brain hemodynamics modulated by blood pressure (Sugimori et al., 1994). Thus, characterizing the arterial-CSF system may help us understand biomechanical modulation of CSF flow.

As an alternative mechanism, recent work has proposed that the autonomic nervous system (ANS) plays an important role in CSF modulation (Picchioni et al., 2022). In particular, in view of the fact that respiration is controlled by the activity of the ANS (Tattersfield & McNicol, 1987), Picchioni et al. used a cued deep-breathing task to transiently elevated CO_2_ and produced an increase in slow-wave CSF pulsations with lag times consistent with a mechanism involving the autonomic nervous system (ANS) (Picchioni et al., 2022). Moreover, respiratory modulation has been shown to produce CSF net displacement in the brain and spinal cord (Liu et al., 2025). As the ANS can directly control CBV, ANS activity can also alter vascular tone, and ANS tone is directly related to baseline CO_2_ (Fukuda et al., 1989; Polosa et al., 1983). Accordingly, the work of Picchioni et al. suggests that there are two distinct mechanisms driving neurofluid dynamics, namely CBV driven by task-based electrocortical activity mediated, and CBV driven by ANS regulation through sympathetic control of respiration and vascular tone. Indeed, given the fact that changes in pulse volume, sympathetic activity, and inspiratory depth are known to co-occur with CSF fluctuations (especially in the neurogenic band) and the K-complexes that are characteristic of N2 sleep (Colrain, 2005; de Zambotti et al., 2018; Halász et al., 2004), ANS regulation may also mediate the sleep-induced CSF pulsations. It is further proposed that external task-driven and internal ANS-regulated CSF pulsation changes are coordinated through the ascending reticular activating system in the brainstem (Benarroch, 2018).

Beyond vascular tone and respiratory modulation, ANS could potentially modulate CSF flow through other mechanisms. It was recently shown that the heart rate (HR) and respiratory rate (RR) are both closely related to CSF dynamics (Z. Yang et al., 2024). Moreover, it was revealed that high-frequency respiration rate in fact correlates low-frequency CSF flow (Vijayakrishnan Nair et al., 2022). This correlation can be observed peaks around 0.02 Hz frequency CSF flow, which overlaps with the frequency peak associated with sympathetic control of blood flow (Kastrup et al., 1989). As of now, it is still unclear whether this overlap is the result of cross-band modulation of respiration or coactivation of the sympathetic nervous system (SNS) in cerebrovascular, cardiac and respiratory systems.

The study of biomechanical and ANS control of CSF flow has led to many answers but also more questions. In this paper, we focus on the following: (1) when baseline CO_2_ is manipulated, does the biomechanical or ANS mechanism dominate the regulation of low-frequency neurofluid fluctuations? (2) can the ANS modulate low-frequency neurofluid dynamics not only through respiration but also cardiac activity? (3) what roles does the frequency of CSF and vascular fluctuations play in the coordination between ANS-related activity and neurofluid dynamics? To address the questions, this work strives to manipulate biomechanics and ANS activity by manipulating baseline CO_2_ level in a group of healthy young adults. Specifically, hypercapnia increases sympathetic activity and hypocapnia increases parasympathetic activity. Therefore, manipulating the capnias baseline may result in the modulation of CSF dynamics via an ANS-mediated pathway in addition to the CBV-mediated pathway. We characterize the contribution of ANS variables (related to respiration and cardiac pulsation) on CSF and vascular fluctuations as measured using rs-fMRI.

## Methods

### Data set

This analysis was conducted on 13 participants at hypercapnic, normocapnic and hypocapnic baselines, with further exclusion of two participants due to the absence of physiological measurements, resulting in 11 participants (25-38 years old; 2 males and 9 females) being included in the study. Participants were recruited through the Baycrest Participants Database, consisting of individuals from the Baycrest and local communities. The study was approved by the research ethics board (REB) of Baycrest, and the experiments were performed with the understanding and written consent of each participant, according to REB guidelines.

### MRI acquisition

Image acquisition was performed with a Siemens TIM Trio 3 Tesla System (Siemens, Erlangen, Germany), which employed 32-channel phased-array head coil reception and body-coil transmission. rs-fMRI data was acquired using a gradient-echo EPI pulse sequence (TR = 380 ms, TE = 30 ms, FA = 40°, 15 slices, 3.44⨉3.44⨉5 mm^3^ with 20% slices gap, 950 volumes). T1-weighted MPRAGE anatomical image was acquired (TR = 2400 ms, TE = 2.43 ms, FOV = 256 mm, TI = 1000 ms, readout bandwidth = 180 Hz/px, voxel size = 1⨉1⨉1 mm^3^). A finger oximeter built into the scanner was used to monitor heart rate, and a pressure-sensitive belt was used to monitor respiration during the rs-fMRI scan. Data sets without coverage of the aqueduct (n=1) and fourth ventricle (n=4) were excluded from subsequent analyses of aqueduct and fourth ventricle signals.

### Gas manipulation

We administered mixtures of O_2_, CO_2_ and medical air using the RespirAct^TM^ breathing circuit (Thornhill Research, Toronto, Canada) for all gas manipulations. With the sequential gas delivery method (Slessarev et al., 2007), the end-tidal partial pressure of O_2_ (PETO_2_) and CO_2_ (PETCO_2_) pressures were targeted by computerized and independent means. This setup allowed us to manipulate the basal PETCO_2_ level of each participant precisely, without altering PETO_2_. We separately targeted a normocapnic baseline (participant’s nature baseline), a hypercapnic baseline, and a hypocapnic baseline, separated by 4 mmHg CO_2_. There was no change in capnia during the recording in order to avoid the possibility of CSF directly being affected by vasodilation caused by the transition between different capnias (Liu et al., 2025; Zimmermann et al., 2023). The RR was self-regulated throughout the respiratory challenges. Hypocapnia is achieved with RespirAct^TM^ primarily through increased breathing depths, and hypercapnia is achieved by increasing CO_2_ levels in the air supply. A pseudo-randomized sequence of capnic conditions was used across different participants, with approximately two minutes between each condition. During the study, breath-by-breath CO_2_ levels were recorded at a rate of 50 Hz using the RespirAct.

### Data preprocessing and parameterization

The FreeSurfer reconstruction was performed on the T1 anatomical data for all participants using FreeSurfer 6.0 (available at: https://surfer.nmr.mgh.harvard.edu). This reconstruction provided tissue segmentation of gray matter, white matter structures, as well as ventricles, which can then be used to delineate regions of interest.

### rs-fMRI data processing

The rs-fMRI data were preprocessed as follows: 1) discard the first 200 volumes, 2) motion correction, 3) motion regression with 6 parameters (3 translations and 3 rotations), 4) demeaning, 5) coregistration with the anatomical image.

### Vascular signals

Vascular regions of interest (ROIs) were defined using the same method as our previous study (Attarpour et al., 2021). That is, arterial and venous maps generated directly from rs-fMRI data from the top 20 percentile of signal-fluctuation amplitudes delineated. In large arteries, slow rs-fMRI signal fluctuations are driven primarily by dynamic magnetic-susceptibility differences driven by dynamic partial-volume effects between the highly oxygenated arterial blood and surrounding tissue (Tong et al., 2019). In large veins, the rs-fMR signal fluctuations are largely attributable to transverse relaxation-rate variations, which, also as shown in our previous work (Attarpour et al., 2021), demonstrate patterns similar to those in large arteries.

### CSF signals

According to previous research, we also calculated a surrogate of CSF velocity by applying the first temporal derivative to the global-mean BOLD signal (GMS) that was detrended and filtered into 0.01-0.1Hz frequency band; for this purpose, we used a zero-delay fourth-order Butterworth filter (Fultz et al., 2019; H.-C. S. Yang et al., 2022). Moreover, rs-fMRI signals in CSF-related ROIs were extracted. The CSF ROIs were derived either through the use of FreeSurfer tissue segmentation (for the lateral ventricle, third ventricle and fourth ventricle) or by manual delineation (for the cerebral aqueduct) and were downsampled to rs-fMRI space for analysis. Fluctuation in rs-fMRI signal from CSF ROIs reflected the variation in ventricle volume (**Fig. 1c**), whereas fluctuation of temporal derivative of global rs-fMRI signal reflected the inflow and outflow of CSF flow (**Fig. 1d**).

**Figure 1.**
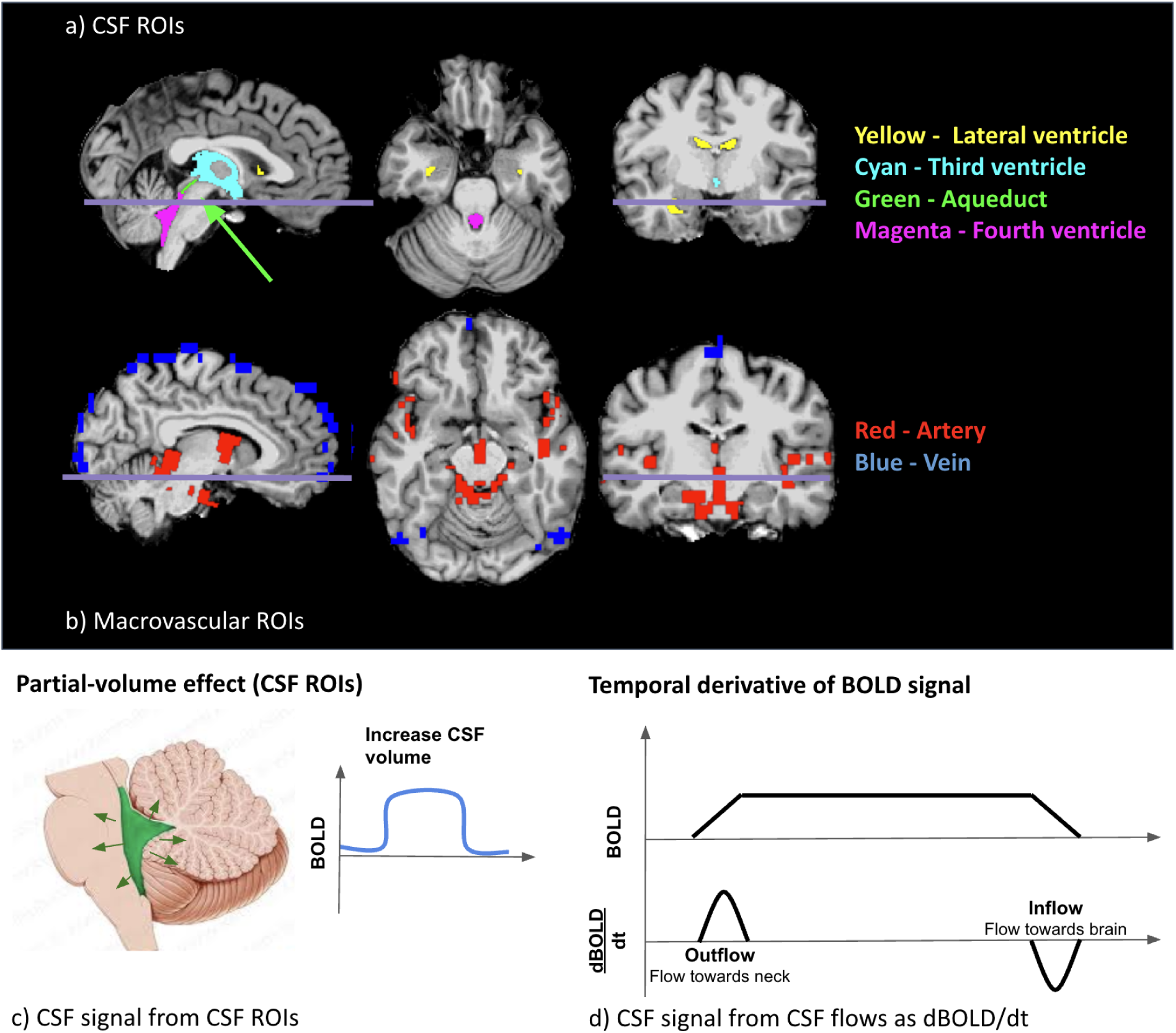
Illustration of the CSF ROI (a) and the vascular ROI (b) from a representative participant. A demonstration of the change in BOLD signal associated with CSF ROIs (d) and flow (c). ROIs were colored as described above, the line indicates the position of the transverse slice. The CSF signal was obtained using two different approaches based on two different mechanisms. BOLD signals increased in CSF signal from CSF ROIs corresponding to volume of CSF ROIs increased (c). A positive BOLD-temporal-derivative CSF flow corresponds to the outflow of CSF (flow towards the neck), while a negative flow corresponds to the inflow of CSF (flow towards the brain) (d).

### Autonomic-related signals

The explanatory variables related to the ANS include: (1) HR; (2) RR; (3) heart rate variability (HRV); (4) respiratory-volume per time (RVT). Both HRV and RVT were estimated based on the method used in the previous study with a window size of 4 seconds (Golestani et al., 2015). Moreover, recent work also found a link between global neuronal activity and the ANS, and accordingly, global-mean gray matter signal (GGMS) was also extracted as a surrogate of global neural activity (Bolt et al., 2022). That is, an increase in GGMS reflects an increase in cortical mean BOLD signal that was found to coordinate with ANS variables such as pulse amplitude and respiratory volume (Tong et al., 2019).

### Frequencies of interest

We identified frequency bands of interest that are consistent with those reported in previous work (Tong et al., 2019), The range of each predetermined band was determined in a data-driven approach and specific to the current group of participants with frequency-domain group-level independent component analysis (ICA) (Calhoun et al., 2001). Specifically, we extracted signals from the neurofluid ROIs of all participants and capnic conditions, and entered them into the ICA. Each band’s range was determined through a virtual search of predetermined three bands among the first 20 independent components. The resulted bands were:

1. Band 1: 0.01-0.14Hz, represents neural activity and vasomotion;
2. Band 2: 0.14-0.56Hz, reflects respiration;
3. Band 3: 0.56-1.31Hz,. reflects cardiac pulsation.

It was found that there is a modulation effect between respiratory measurements and low-frequency CSF flow (Vijayakrishnan Nair et al., 2022). Thus, we divided the BOLD signal in Band 1 into infraslow bands. Based on laser Doppler flowmetry, unstimulated blood flow in Band 1 exhibits 3 spectral peaks, namely:

1. The endogenic peaks (Band IS1): 0.001-0.02 Hz, represents an endogenic (metabolic) band related to the rhythmic regulation of vascular resistance to the blood flow triggered by variations in blood metabolic substrate concentration (Bracic & Stefanovska, 1998);
2. The neurogenic peak (Band IS2): 0.02-0.04 Hz, may result from sympathetic neuronal (SNS) activity (Kastrup et al., 1989);
3. The myogenic peak (Band IS3): 0.06-0.14 Hz, which is associated with blood-pressure regulation (Johnson, 1991), and within the range of low-frequency HRV (Akselrod et al., 1981).

Following the formulation in (Zhong & Chen, 2022), we further calculated the normalized power (power normalized by signal temporal mean) and the central frequency of each frequency range is computed as the frequency at the “centre of mass” of the spectrum, as follows:

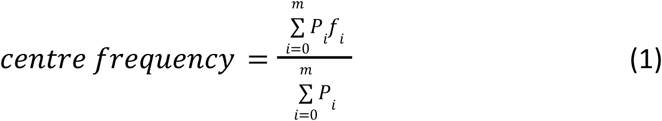

where *P* represents power, and *f* frequency, and (*i* = 0,…,*m*) is the frequency index in the Fourier domain (*m* corresponds to the index of the maximum frequency).

## Statistical analysis

### Power spectra across capnic conditions

Power spectra were calculated using Welch’s method (Hamming window, 41 sample overlaps) and averaged across all participants for each capnic conditions and each ROI.

### Power and central frequency differences across capnic conditions

To demonstrate the difference between power and central frequency for three different capnics, effect sizes were calculated for each band using Glass’s estimator (D) for each frequency band (J. Cohen, 1977; Hedges, 1981). The effects were further classified according to the standard set by the previous study (Sawilowsky, 2009) as follows: very small (D < 0.2), small (0.2 < D < 0.5), medium (0.5 < D < 0.8), large (0.8 < D < 1.2) and very large (D > 1.2).

### Association between physiological factors and power and central frequency

An analysis of linear mixed effects was conducted to investigate the association between each physiological factor and the BOLD signal power and central frequency for each frequency band. The input parameters were all demeaned and normalized by their respective inter-participant standard deviation. The variable “ID” was included in the model as a random variable in order to account for repeated measures within each participant (identified by a participant ID number) (**Eq. 2**). Thus, the inter-participant variation is not of interest, as we are modeling the inter-capnic effects as the outcome measure. The model is illustrated in **Table 1**. Note that in Band 3, since HRV (Holland & Aboy, 2009) and RVT (Birn et al., 2006) are unlikely to extend into such a high-frequency band, only lower-frequency band metrics were incorporated into the linear mixed-effects model (*RVT*(*t*) power and frequency from Band 1 and *HRV*(*t*) power and frequency from Bands 1 and 2 were used as separate independent variables in the LMEs for all rs-fMRI signal bands). The 95% confidence interval of each coefficient in the model is determined by bootstrapping (10,000 iterations of resampling with replacement), and each coefficient is deemed significant only if its confidence interval does not span zero.

**Table 1.** Details on the linear mixed effect models. Y represents the dependent variable and X the independent variable. A separate LME is constructed for each variable under the Y columns, using all variables under X columns (where power and frequency correspond to variables under the Y column).

| $Y \sim 1 + (1 \vert \text{Participant ID}) + \sum X$ (Eq. 2) | | | |
| --- | --- | --- | --- |
| Y (Neurofluid Signal) |  | X (Physiological Recordings) |  |
| Power | Frequency | Power | Frequency |
| CSF Flow | CSF Flow | Respiratory Rate (RR) |  |
| Artery | Artery | Heart Rate (HR) |  |
| Vein | Vein | PETCO<sub>2</sub> |  |
| Lateral Ventricle | Lateral Ventricle | Heart Rate Variability (P(HRV(t))) | Heart Rate Variability (f(HRV(t))) |
| Third Ventricle | Third Ventricle | Respiratory-Volume per Time (P(RVT(t))) | Respiratory-Volume per Time (f(RVT(t))) |
| Aqueduct | Aqueduct | **•The 95% confidence interval with 10000 bootstrapping**<br>**•Significant only if its confidence interval does not span zero** |  |
| Fourth Ventricle | Fourth Ventricle |  |  |

### Temporal characterization of ANS coordination: respiratory and cardiac response functions across capnias

The RRF was estimated to understand the mechanism by which the ANS affects BOLD signal dynamics through *RVT(t)*. For each participant, the mean signal from each neurofluid ROI as well as the GGMS and CSF velocity time series were deconvolved by the corresponding *RVT(t)* using the Laguerre expansion (Prokopiou et al., 2018; Shams et al., 2022) for all macrovascular and CSF ROIs as well as GGMS listed above across three capnias. The cardiac response function (CRF) was also estimated to link the ANS-driven BOLD dynamics through *HRV(t)*. The Wilcoxon signed rank test (p<0.05) was used to determine whether the peak height, lag, and full-width-half-maximum (FWHM) differed between pairs of capnias for the first and second peak as well as the centre frequency (**eq. 1**) for CRF and RRF.

### Temporal characterization of vascular-tone response: arterial response function across capnias

To further separate the effects of ANS-induced effects on from biomechanical effects on CSF dynamics across capnias, the arterial transform function (ATF) was estimated using the same approach as CRF and RRF estimation, by substituting the *RVT(t)* and *HRV(t)* time courses with the arterial BOLD signal (See *Respiratory and cardiac response function across capnias* section). The Wilcoxon rank sum test (p<0.05) was used to determine whether the peak height, lag, and FWHM differed between pairs of capnias for both the first peak as for *ATF(t)*.

### Shared information between BOLD and respiratory/cardiac variability time series

To quantify the dynamic interaction between BOLD and the ANS parameters *HRV(t)* and *RVT(t)*, we calculated the mutual information between these pairs of variables. This permitted us to assess the interactions unbiased by the accuracies of the response functions. Mutual information was calculated using a MATLAB toolbox (Feed et al. 2015) with the window width (k-nearest-neighbours) chosen to be consistent with the 0.1 Hz bandwidth of the rs-fMRI signal (26 temporal samples). The mutual information index was normalized by the geometric mean. The Wilcoxon signed rank test with a threshold of 0.05 was performed to determine whether there were differences between capnias.

## Results

In **Table 2**, we illustrated the physiological metrics across the 2 capnic conditions, including PETCO_2_, heart rate (HR), and respiratory rate (RR). As expected, hypercapnia was associated with the highest PETCO_2_ levels, followed by normocapnia, with hypocapnia exhibiting the lowest levels. The highest HR was associated with hypercapnia, followed by hypocapnia, with the lowest HR associated with normocapnia. There was an opposite trend in the RR, where normocapnic RR was the highest, followed by hypercapnic and lastly hypocapnic RR. Significant differences across capnic levels were found for PETCO_2_ but not for the HR and RR in terms of group mean (not equivalent to pairwise differences). Additionally, **Figure 2** shows power spectra for BOLD signals from vascular and CSF ROIs as well as HRV and RVT.

**Table 2.** Physiological measurements (mean and standard deviation). P-values were computed by one-way analysis of variance (ANOVA).

|  | Hypercapnia | Normocapnia | Hypocapnia | P-value |
| --- | --- | --- | --- | --- |
| PETCO<sub>2</sub> (mmHg) | 42.91±2.47 | 39.47±1.93 | 36.44±1.60 | 1.41e-07 |
| Heart rate (Hz) | 1.28±0.32 | 1.15±0.18 | 1.26±0.31 | 0.54 |
| Respiratory rate (Hz) | 0.23±0.07 | 0.23±0.06 | 0.22±0.06 | 0.93 |

**Fig. 2.**
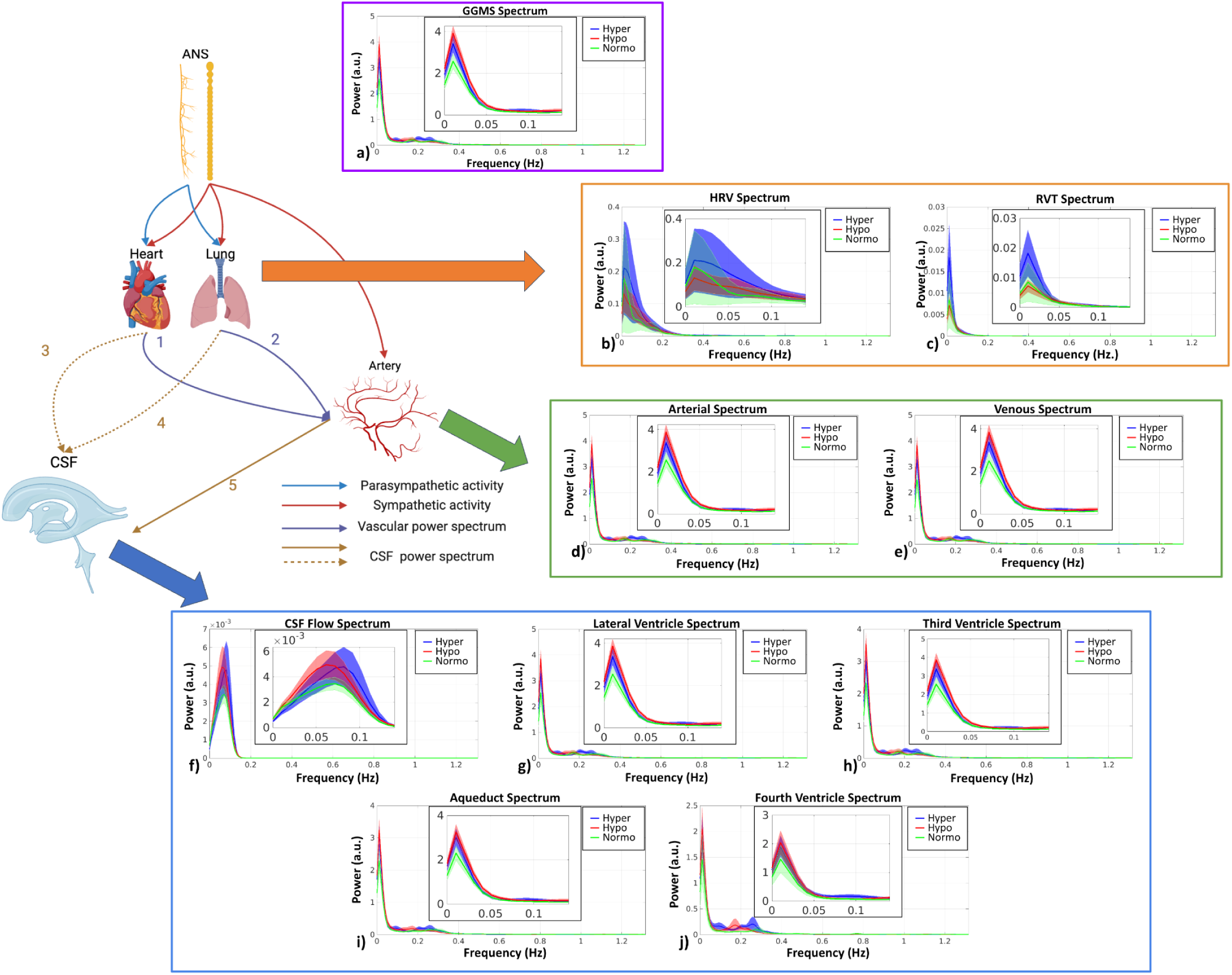
BOLD signal power spectra in all ROIs. The schematic illustrates the pathways under investigation that link the ANS to CSF dynamics. The heart and lung both receive parasympathetic (blue arrow) and sympathetic (red arrow) input, whereas the arterial vasculature only receives sympathetic input (red arrow). The lung and the heart may have a direct impact on arterial BOLD (purple arrows labeled ‘1’ and ‘2’), which in turn directly drive. CSF flow (brown arrow labeled ‘5’), as well as indirectly by heart (blue dashed arrow labeled ‘3’) and lung (blue dashed arrow labeled ‘4’) activity. All power spectra were calculated after time-course normalization by the temporal mean. The shaded area represents standard error. The zoomed-in versions (Band 1) of the spectra are shown as insets. (Figure partially based on biorender: https://www.biorender.com/).

### Vascular and CSF oscillations across different capnias

Signal metrics were compared across capnic conditions, and all associated results are shown as Cohen’s D values, as summarized quantitatively in **Fig. S3** as well as verbally in **Table S1**. When comparing hypocapnia to normocapnia, the BOLD signal in vascular and CSF ROIs behaved in very similar ways, and the same is true when comparing hypercapnia to normocapnia. Relative to normocapnia, both hypocapnia and hypercapnia are associated with increased BOLD signal power and shifts in BOLD signal frequency in a band-dependent manner. The behaviour of CSF velocity, only defined for frequencies up to 0.1 Hz, also mimics that of the BOLD signals in the vascular and CSF ROIs. Moreover, GGMS signal metrics behaved very similarly to those from vascular ROIs.

### Drivers of vascular and CSF oscillations across capnic conditions

We see obvious differences across the infraslow frequency bands and non-infraslow frequency bands (**Figure 3**). Interestingly, basal CO_2_ was not a strong driver of the differences across capnic conditions for all frequency bands. Below are the details of the effect for each band:

● Band IS1: Positive associations between frequency and RR in the artery, vein, lateral ventricle, third ventricle, and GGMS.
● Band IS2: Negative association between frequency and HR in veins; negative association between CSF flow frequency and HRV (f(*HRV*(*t*)).
● Band 1: Negative association between RR and frequency in arteries, veins, the lateral ventricle and third ventricle; positive association between frequency and HRV (f(*HRV*(*t*)) in the GGMS.
● Band 2: Positive association between HRV (f(*HRV*(*t*)) and GGMS frequency

**Figure 3.**
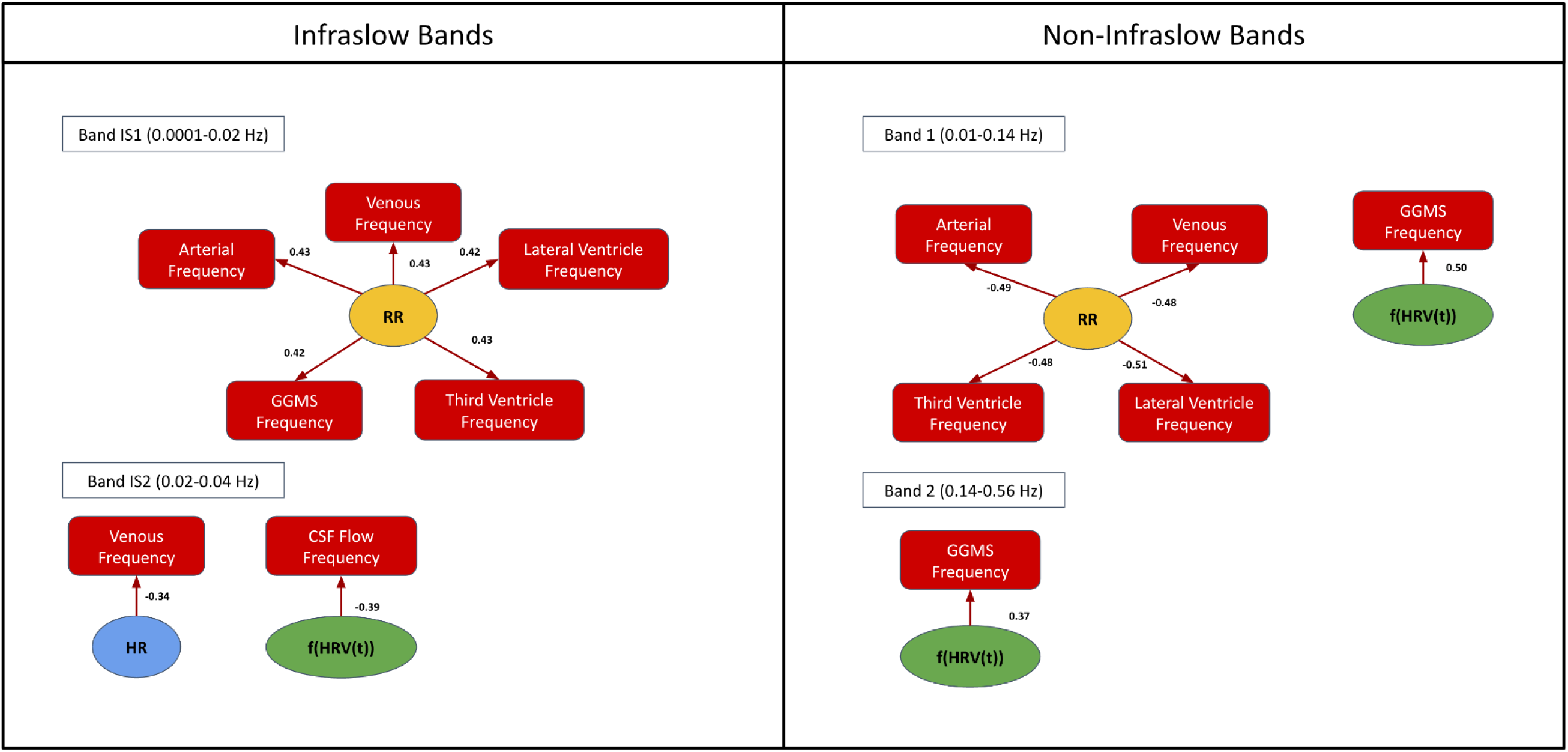
Factors linked to CSF signal fluctuations in the infraslow bands IS1-IS3 (left) and band 1-3 (right). Only significant results are shown (p<0.05 with the bootstrapped test). The arrows indicate the significant association. The numbers displayed next to the arrows indicate the effect size. HR: heart rate; RR: respiratory rate; HRV: heart-rate variability.

It is also worth noting that extra-cranial sources in upper frequencies (Bands 2 and 3) do not significantly modulate neurofluid dynamics in our modeling framework.

### RRF and CRF for different capnic conditions

In this work, we use the RRF and CRF to characterize the time-dependent coordination between the ANS and CSF/vascular signals. Based on the fact that RR is the most predominant mediator of the relationship between capnic condition and BOLD signal changes in neurofluid ROIs, we further estimated the RRF linking respiratory variability to the BOLD data. As shown in **Figure 4**, there were distinguishable differences in the shapes of the RRF across three capnias. Further statistical analysis revealed that hypercapnia showed a higher first FWHM than normocapnia for all ROIs and GGMS, but not for CSF flow. The hypercapnia also showed higher first peak lags than hypocapnia for lateral and third ventricles.

**Figure 4.**
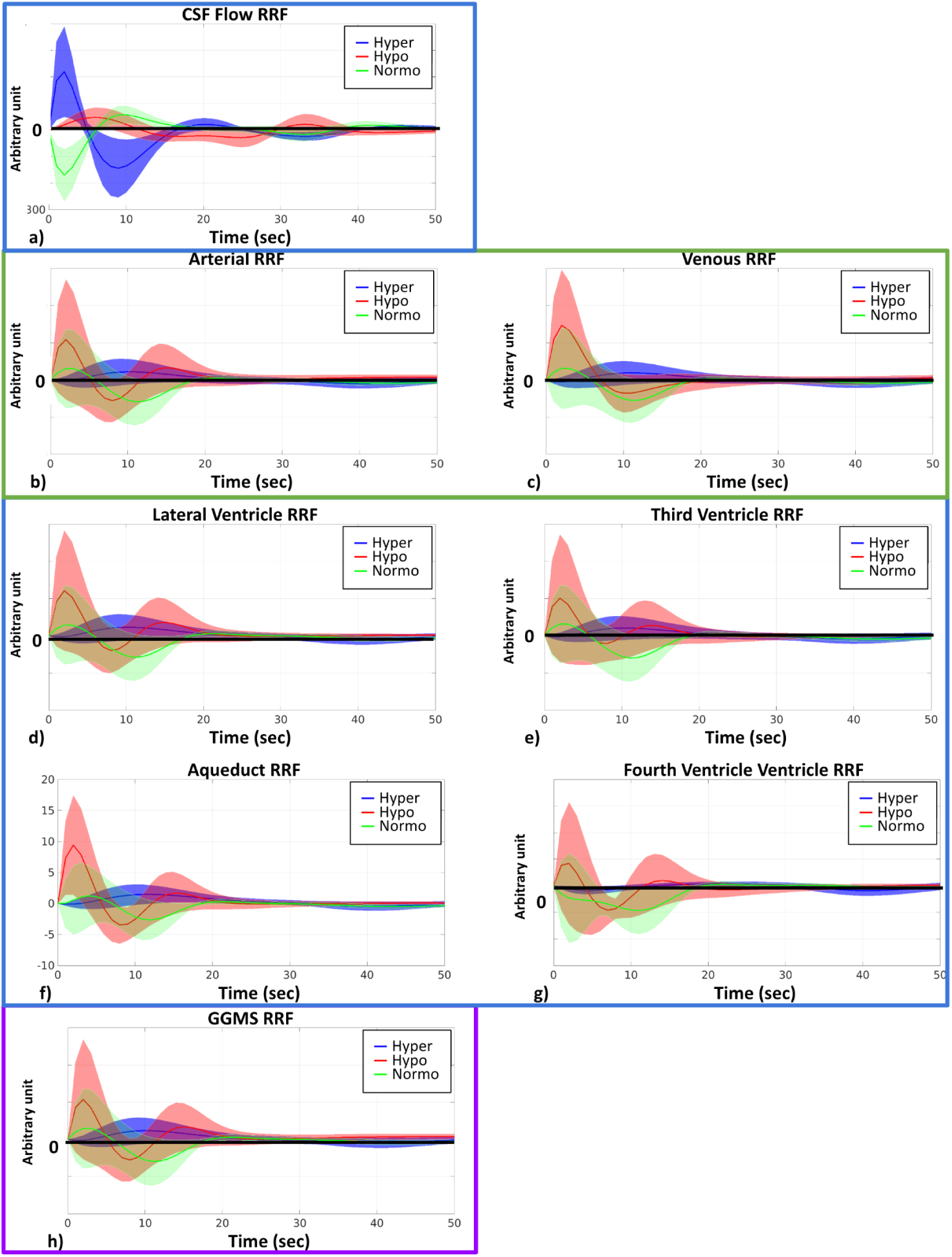
RRF across different capnias. All plots are shown as averages across participants, and the shaded area represents the standard error. The y-axis of plots is in arbitrary units. Hyper (blue): hypercapnia; hypo (red): hypocapnia; normo (green): normocapnia.

As shown in **Figure 5**, there were distinguishable differences in the shapes of the CRF across three capnias. Further statistical analysis revealed that the first peak intensity for the CRF for hypocapnia was significantly higher than normocapnia for CSF flow. Moreover, normocapnia showed higher second peak intensity than hypocapnia for CSF flow.

**Figure 5.**
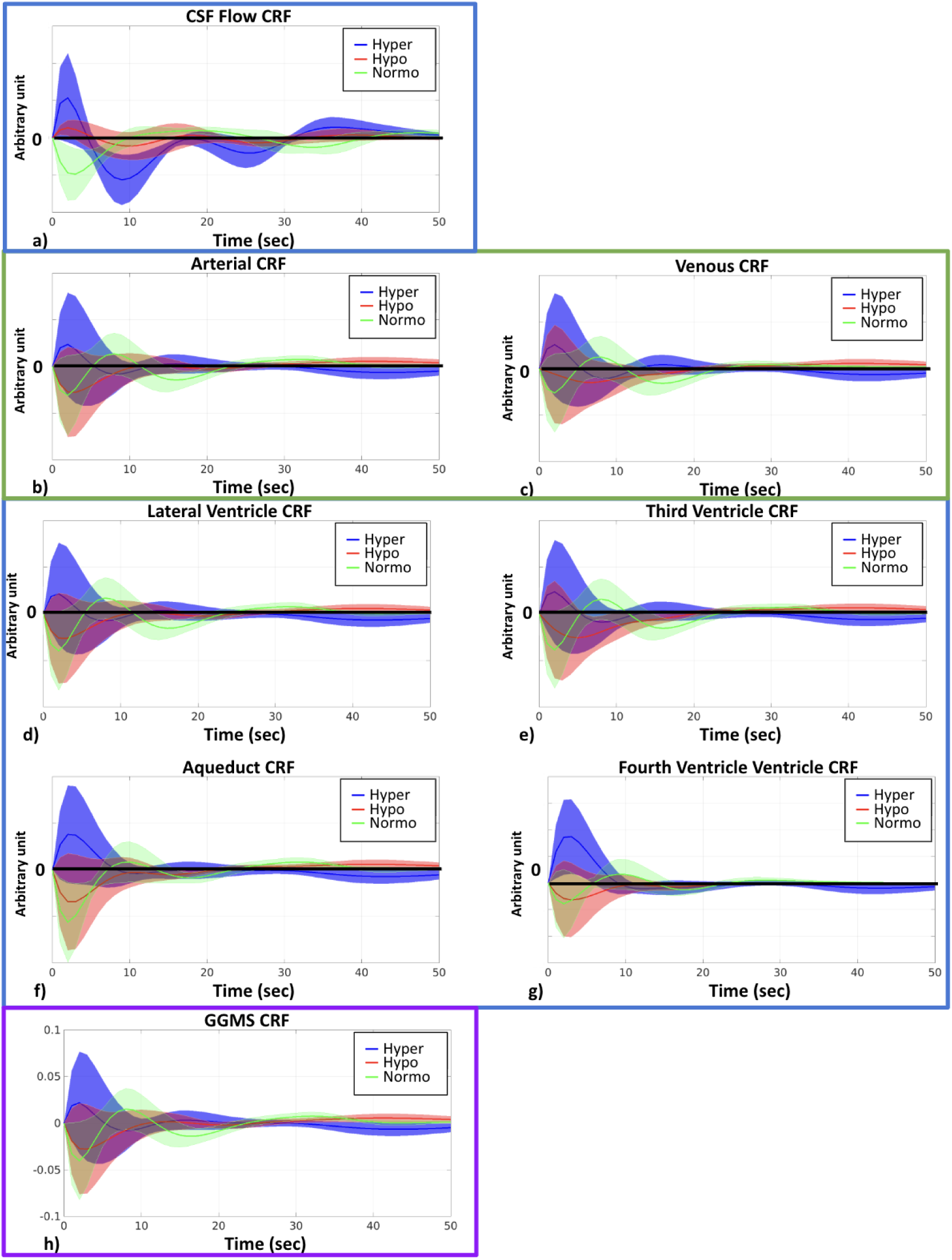
CRF across different capnias. All plots are shown as averages across participants, and the shaded area represents standard error. The y-axis of plots is in arbitrary units. Hyper (blue): hypercapnia; hypo (red): hypocapnia; normo (green): normocapnia.

### Arterial-transfer function (ATF) for different capnic conditions

As the CSF flow dynamic in the resting state (after the capnic effect reach a steady state) originates primarily from vascular activity, especially arteries, given their ability to dilate and contract, ATF was estimated in order to better understand how ANS-induced vascular tone differences impact CSF dynamics. The ATF quantitatively describes how arterial fluctuations are converted to CSF flow fluctuations, a process which is directly influenced by tissue elasticity and vascular tone, as described earlier. As outlined in the biophysical model (Zhong et al., 2024), the increase in arterial signal is mainly due to the contraction of the arteries (less R ^’^ decay caused by arterial-derived local-field inhomogeneity). As shown in **Figure 6**, ATFs from different CSF ROIs and CSF flow behave very similarly at different capnias. Further statistical analysis revealed there is no significant difference in the ATF parameters across different capnias, although qualitative differences exist in the plots for CSF ROIs.

**Figure 6.**
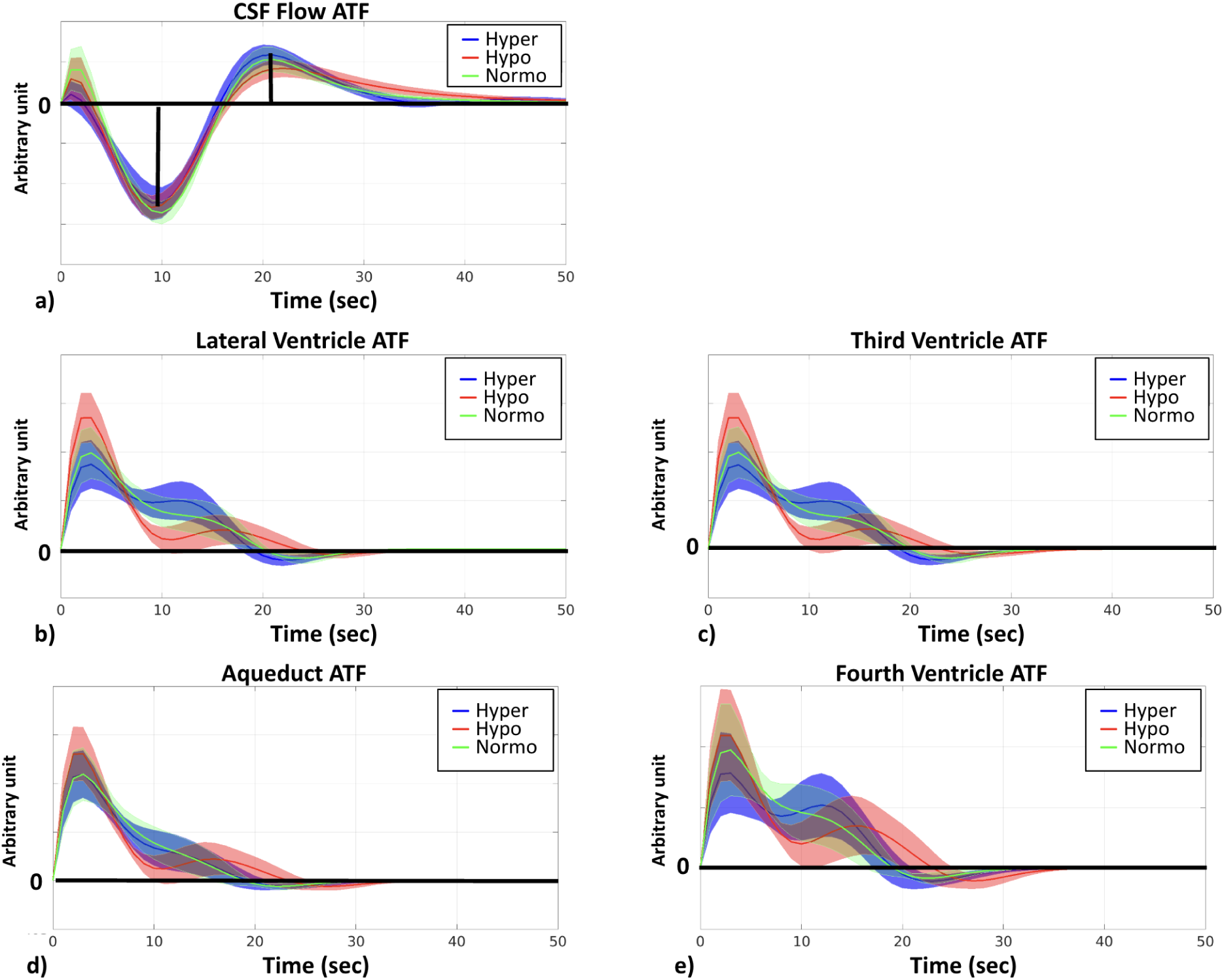
ATF across different capnias. All plots are shown as averages across participants, and the shaded area represents the standard error. The y-axis of plots is in arbitrary units. Hyper (blue): hypercapnia; hypo (red): hypocapnia; normo (green): normocapnia.

Given that the ATF did not differ significantly across capnias, whereas the RRF and CRF did, and given that we mainly uncovered differences in signal frequency across capnias (**Fig. 7a-d**) that are mediated by *HRV*(*t*) and *RVT*(*t*) frequencies (**Fig. 3**), we wanted to clarify if these frequency differences in **Figure 7a-d** are reflected in the RRF and CRF frequencies. As shown in **Figure. 7e-l**, frequency differences across capnias were identified in the RRF estimates associated with all signals, but not in the CRF estimates. This suggests that despite *HRV*(*t*) frequency (and not *RVT*(*t*) frequency) being the major ANS-related mediator of neurofluid signal frequency differences across capnias, the frequency governing the propagation of HRV variations to neurofluids variations remains unaffected by capnia, whereas the frequency governing the propagation of RVT variations to neurofluids variations is capnia-dependent. The frequency differences for CSF ROIs can be found in Supplementary Materials (**Figure S4).**

**Figure 7.**
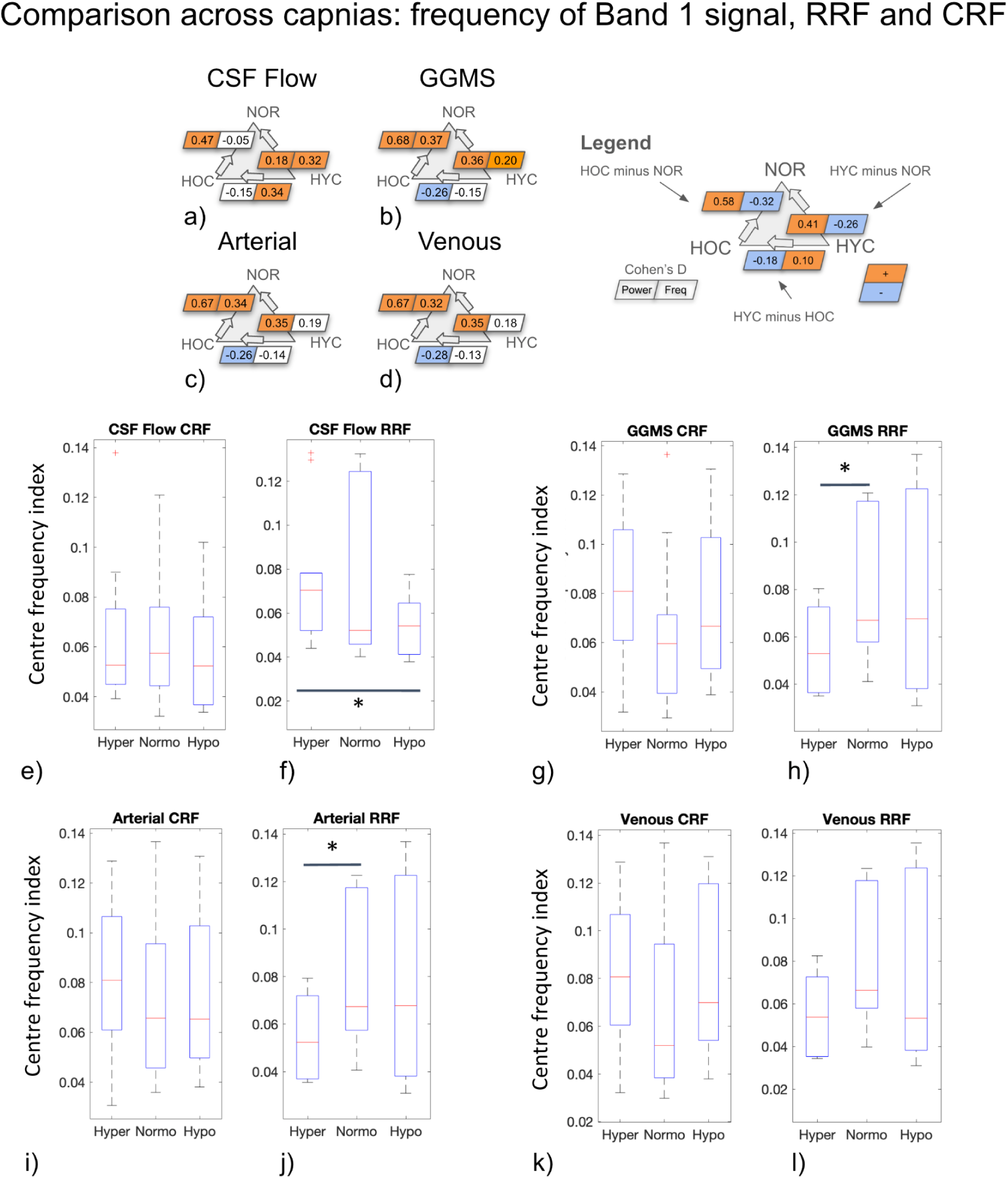
Summarized in (a-d) are the differences in power and centre-frequency index of Band 1 signal fluctuations in CSF flow, GGMS signal, the rs-fMRI signal in the arterial and venous ROIs, respectively. Correspondingly, (e-l) summarize the centre-frequency indices of RRF and CRF calculated for CSF flow, the GGMS signal, the rs-fMRI signals in the arterial and venous. Asterisks indicate significant differences based on the Wilcoxon signed test (p < 0.05). HYC: hypercapnia; HOC: hypocapnia: NOR: normocapnia.

### Interaction between BOLD and HRV/RVT time series

To further understand the sizes of contribution of respiratory and cardiac variability to neurofluid dynamics, we compared mutual information measures relating these latters to neurofluid waveforms. Compared to *HRV*(*t*), *RVT*(*t*) showed higher amounts of mutual information with CSF flow (**Fig. 8**). Nevertheless, the mutual information analysis did not reveal significant differences across capnic conditions. There was, however, a trend of increasing mutual information from hypocapnia to hypercapnia. Details for all ROIs can be found in Supplementary Materials (**Figure S2)**.

**Figure 8.**
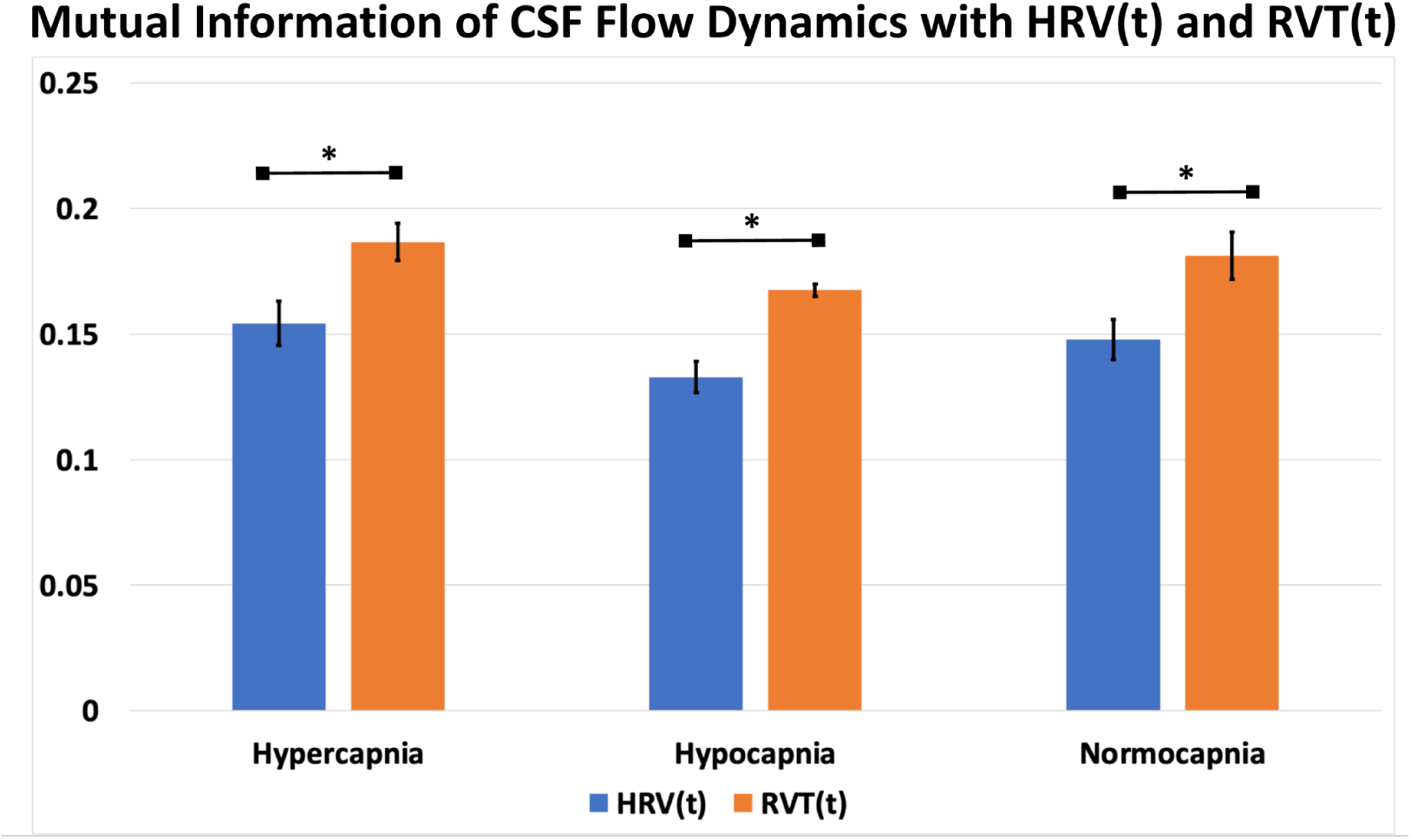
Comparison of mutual information of CSF flow dynamics with *HRV(t)* and *RVT(t)*. Blue: *HRV(t)*; orange: *RVT(t)*. The error bars represent the standard error across participants and the asterisk indicates significant differences.

## Discussion

It is increasingly established that fMRI-based CSF fluctuation measurements reflect variations in CBV. Dynamic properties of these CBV variations can in turn be modulated biomechanically by vascular tone and neuronally by the ANS. However, it remains unclear to what extent each pathway contributes under circumstances of altered vascular tone and altered ANS tone. In this work, we dissected these different mechanisms by measuring neurofluid dynamics as reflected by rs-fMRI during altered steady-state CO_2_, i.e. during hypercapnia, normocapnia and hypocapnic baselines. Our main findings are:

1. While CO_2_ can modulate CBV, the manner biomechanical modulation remains unaltered across capnic conditions, and do not drive the variations in neurfluid dynamics across capnias;
2. In addition to respiration, as previously reported, cardiac pulsation also independently drives neurofluid flow as an indication of the ANS pathway of control.
3. Altered CO_2_ alters neurofluid dynamics primarily through the frequency (instead of the amplitude) of heart-rate and respiratory-volume variability.

In this work, we prioritize the derivative-based CSF flow velocity time course for representing CSF dynamics, as it is distinct from the signals from the CSF ROIs, which are in turn more similar to those of arterial and venous ROIs (**Fig. 4** and 5). This is in agreement with our previous findings (Attarpour et al., 2021) as well as with based on cine phase contrast (ElSankari et al., 2013) and highlights the differences in the contrast mechanisms between CSF flow and CSF-BOLD (**Figure 1**).

### The biomechanical perspective

While CO_2_ is a widely used method to alter vascular tone and hemodynamics (Ainslie & Duffin, 2009; E. R. Cohen et al., 2002), which are in turn tightly linked to CSF dynamics, the mechanisms through which basal CO_2_ influences CSF dynamics are less well documented. Specific to this work, it is not entirely clear to what extent this was due solely to the biomechanical effect of basal CO_2_ on CVR, and due to the neuromodulatory effects of CO_2_. Vascular tone is conventionally defined as the baseline level of constriction in a blood vessel relative to its maximally dilated state. However, in the context of fMRI signals, the ability to fluctuate depends not only on dilatory capacity but also on constrictive capacity. Thus, in the context of rs-fMRI based neurofluid fluctuation measurements, a more neutral vascular tone (i.e. during normocapnia) stands to be biomechanically associated with the highest level of signal fluctuations, as was shown in our previous work (Halani et al., 2015). This is consistent with previous work by Biswal et al., who uncovered a suppression of 0.08 Hz band rs-fMRI signal fluctuations during 5% hypercapnia, which was attributed to the reduced sensitivity of hemodynamic modulation during a vasodilated state (Biswal et al., 1997). These observations can in part explain our own observations of significant positive correlations between rs-fMRI signal fluctuation amplitude and CVR, with CVR being diminished at hyper- and hypocapnia compared to normocapnia (Golestani et al., 2016). Moreover, previous work by Cohen et al. (E. R. Cohen et al., 2002) found that the BOLD peak height was inversely correlated with basal CO_2_, while the BOLD response width was directly correlated with basal CO_2_.

As CVR directly reflects CBV fluctuation amplitude, the biomechanical hypothesis would predict that the higher CVR at normocapnia be associated with higher CSF fluctuations than at hyper- or hypocapnia, and that the hypocapnic baseline be associated with faster CSF fluctuations. However, this was not what we observed. Instead, we found CSF as well as vascular fluctuation amplitude and frequency to both be higher at hyper- and hypocapnia (**Fig. 7**). Moreover, given the ATF is unchanged across capnias (**Fig. 6**), implying that the ability of arterial pulsations to drive CSF pulsations is independent of vascular tone and CVR, the biomechanical mechanism is unlikely dominate the differences in the dynamics of CSF across capnias.

### The role of the ANS: respiration and cardiac pulsation

There are complex interactions between the ANS and neurofluid flow (Halani et al., 2015; Picchioni et al., 2022), as there are between basal CO_2_ and ANS tone. Respiratory variability is a key metric associated with ANS function (Chang et al., 2013; Lane et al., 2009; Lehrer & Gevirtz, 2014; Napadow et al., 2008). As part of the sympathetic reflex, levels of CO_2_ and O_2_ in the blood are monitored by chemoreceptors in the brainstem and carotid bodies. Increased CO_2_ or decreased O_2_ levels trigger the sympathetic nervous system (SNS) to increase RR and HR to maintain proper gas exchange (Davis et al., 1977), thereby modulating RVT and leading to reduced HRV. Changes in RR, HR, RVT and HRV can also alter not only the signal fluctuation amplitude but also the frequencies of their corresponding fMRI signal components, and result in altered frequency content in specific bands of rs-fMRI signal (Birn et al., 2006; Chang et al., 2009). In our analysis, RR were found to significantly correlate with rs-fMRI signal fluctuation frequency in neurofluid ROIs in the low-frequency band (Band 1 and Band IS1) as well as GGMS (Band IS1) frequency. HR and HRV were also found to be significantly associated with venous (Band IS2) frequency, GGMS (Band 1-2) frequency and CSF flow (Band IS2) frequency. In this study, we found associations primarily through frequency coupling. This phenomenon suggests that frequency transfer might be more linear and less distort than power transfer along the neurofluid system (from heart or lungs to vasculature and CSF flow), particularly in light of the complicated turbulent flow of blood (Saqr et al., 2020) and CSF (Atsumi et al., 2020). A better understanding of the power and frequency couplings adds a new dimension to previous studies that mainly focused on the temporal pattern of CSF dynamics (Picchioni et al., 2022; Z. Yang et al., 2024).

In terms of the temporal analysis, hypercapnia is associated with a very slowly varying RRF (**Fig. 4**), which is reminiscent of the slowly-varying BOLD response that Cohen et al. observed during a hypercapnic baseline (E. R. Cohen et al., 2002). In contrast to RVT, HRV mainly modulates neurofluid oscillation through an amplitude modulation, in which hypocapnia results in a higher first peak and a lower second peak (**Fig. 5**). Moreover, the flip from positive (in hypocapnia) to negative (in normocapnia) for the first peak (**Fig. 5**) indicates that the direction of flow associations with HRV is also different across capnias. Furthermore, our mutual information indicates that RVT exhibits higher mutual information with CSF flow than does HRV, indicating the respiratory system is the main determinant of ANS-CSF modulation. Thus, not only do our results show that the manner in which both HRV and RVT modulate neurfluid dynamics varies by capnia, these results clarify the manner in which neurofluid dynamic in the low-frequency band differs across capnic conditions. A broader implication could be that different individuals can be found on different points on the capnic spectrum, basal capnia is also an important contributor to inter-participant and inter-session variability in RRF, as well as in neurofluid flow patterns.

### The role of ANS control of vascular tone

Similar to the heart and lung, the microvasculature also receives input from the ANS, more specifically the sympathetic nervous system (**Fig. 9**). In fact, previous findings by Peebles et al. supported the regulation of CVR by the sympathetic nervous system through the alpha1-adrenoreceptors (Peebles et al., 2012). Elevated SNS tone triggers the release of norepinephrine from sympathetic nerve endings near blood vessels. Norepinephrine binds to alpha-adrenergic receptors on smooth muscle cells in the blood vessel walls, causing them to contract. This vasoconstriction also leads to increased MAP (Smith & Maani, 2023) and reduced blood flow (CBF) as well as CBV, increasing vascular tone (Jordan et al., 2000). In the brain, this vasoconstriction may also divert blood towards the heart and limbs, except for in regions related to the stress response (ter Laan et al., 2013). As we did not monitor blood pressure (MAP) during our capnic conditions, we cannot comment on the role of the vagal reflex in inducing RR and HR changes. A recent study suggests a negative correlation between SNS activity and peripheral vasomotion in anticipation of pain stimuli (Z. Xu et al., 2024). Thus, the pathways of ANS influence can be summarized as: increased CO_2_ → increased SNS tone → reduced CBF and CBV → reduced vasomotion. However, this is not what we observed, as we observed higher neurofluid oscillation amplitude at higher CO_2_ levels. Thus, the interplay between the ANS and neurofluid dynamics does not appear to be dominated by ANS control of vascular tone. Moreover, to address an earlier point regarding the existence of coactivation of the SNS cerebrovascular, cardiac and respiratory systems, as the ATS did not vary across capnias as the RRF and CRF did, our data does not support coactivation.

**Figure 9.**
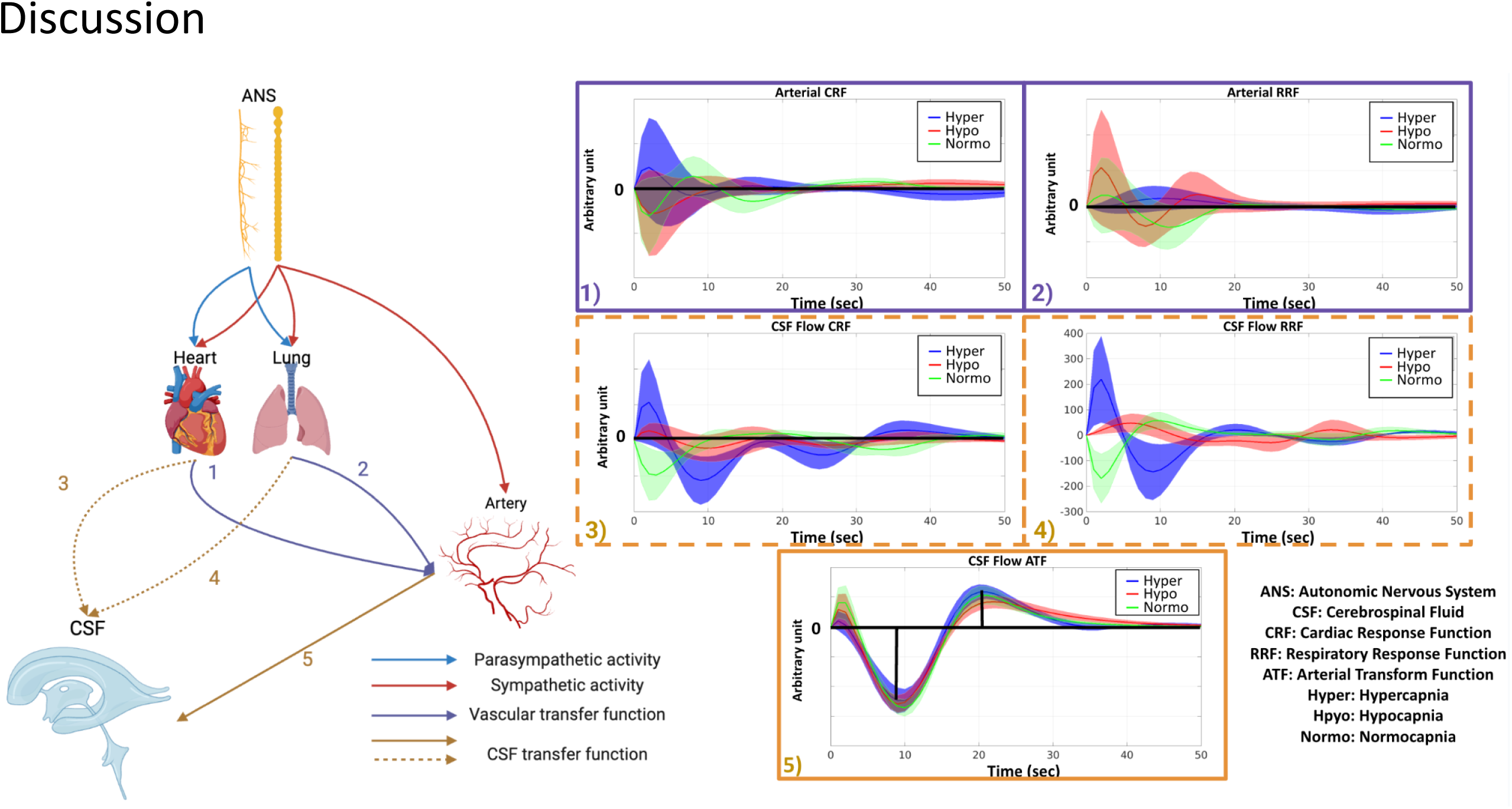
An overview of the pathways that drive the flow of CSF by the ANS. The lung and heart both receive parasympathetic (blue arrow) and sympathetic (red arrow) input, whereas the arterial vasculature only receives sympathetic input (red arrow). The lung and the heart may have a direct impact on arterial BOLD (purple arrows 1 and 2), which is modulated by arterial CRF (1) and RRF (2). The CSF flow is directly driven by vascular pulsation (brown arrow 5), modulated by CSF flow ATF (5), as well as indirectly by heart (blue dashed arrow 3) and lung (blue dashed arrow 4) activity, which are modulated by CSF flow CRF (3) and RRF (4). (Figure partially based on biorender: https://www.biorender.com/). Hyper (blue): hypercapnia; hypo (red): hypocapnia; normo (green): normocapnia.

### Beyond the ANS

Aside from the ANS-mediated modulation of CSF dynamics, baseline CO_2_ may modulate CSF dynamics through other potential pathways. We will discuss these potential pathways in more detail below.

#### CO_2_, heart and respiratory rate

Baseline CO_2_ may alter the heart and respiratory rate directly through stimulating chemoreceptors (Kronenberg & Drage, 1973). While the group comparisons across capnic conditions did not reveal a significant difference in HR or RR, on an individual level, HR and RR still differed across conditions (**Figure S1** in Supplementary Materials). At a group level, while not significant, HR is higher at both hyper- and hypocapnia. As well, RR is highest at hypercapnia and lowest at hypocapnia. As our analyses are band-specific, frequency shifts due to such changes can result in seemingly counterintuitive observations. For example, an upward shift of the HR-related BOLD signal peak can result in the aliasing of the HR peak into Band 1. However, based on our HR measurements, this only applies to 3 of the participants (**Fig. S1** in Supplementary Materials). Accordingly, based on the limited group effects observed in our experiment, baseline CO_2_ modulation of neurofluid by directly affecting HR and RR is not comparable to the effect of ANS-mediated modulation.

#### CO_2_ and neuronal activity

An alternate or complementary mechanism for the effect of CO_2_ on the rs-fMRI signal is through the effect of CO_2_ on neural activity. Elevated CO_2_ has been associated with suppressed steady-state amplitude of the band-limited EEG Hilbert envelope amplitude in multiple EEG and MEG bands (Driver et al., 2016; F. Xu et al., 2011). By the same token, reduced CO_2_ should be associated with enhanced EEG amplitude. In this regard, previous work suggests that neuronal activity variations can influence hemodynamics, which in turn influence CSF fluctuations as measured using BOLD rs-fMRI. Fultz et al. (Fultz et al., 2019) found that the global grey-matter BOLD signal was coupled with the CSF BOLD signal from the fourth ventricle, which was in turn coupled with low-frequency delta activity. Following the work of Fultz et al., (Fultz et al., 2019) if hypercapnia and hypocapnia act to suppress and enhance synchronized neuronal activity fluctuations, respectively, they can also enhance and suppress the CSF signal fluctuations, respectively.

In this experiment, we used GGMS as a proxy for global neural activity. The GGMS originates from all grey matter and has also been associated with global activity, particularly within the low-frequency range (Band 1) (Bolt et al., 2022). Here, we confirm that what we observed in arterial, venous and ventricular ROIs can be mirrored in the GGMS. However, the origins of the GGMS are far from clear. For instance, neural activity has been shown to entrain regional vasomotion (Drew et al., 2020; Mateo et al., 2017), while subdural vascular oscillations in the cranial bone has also been found to closely resemble the GGMS (Huber et al., n.d.) suggesting that the GGMS also contains systemic vascular oscillations (Tong et al., 2013). It is therefore impossible to rule out the possibility that the baseline CO_2_ may alter CSF dynamics through a non-ANS neuronal-mediated pathway. However, since the neuronal hypothesis would suggest that hypercapnia and hypocapnia should be associated with opposite effects on CSF fluctuations relative to normocapnia, which was not the case, as seen in **Figure S3** and **Tables S1** and **S2**, neuronal activity changes are not the main driver of the CSF-flow differences across capnias.

### Implication of CSF fluctuation amplitude and frequency

While the frequency of CSF flow fluctuations and BOLD signal fluctuations can both increase with respiratory rate (RR) as part of hypercapnia and hypocapnia (Vinje et al., 2019), in our data, RR was maintained across all capnic conditions (**Table 1**). Moreover, RR does not necessarily correlate with low-frequency variations in RR. Thus, a more interesting question is what underlies the link between frequencies of *HRV*(*t*), *RVT*(*t*) and neurofluid signal frequency. As shown in **Figure 7**, the frequency of RRF, which links *RVT*(*t*) and neurofluid signals, varies with capnia. Thus, in this low-frequency range, altered CO_2_ level results in a faster or slower translation from respiration to neurofluid flow.

Current literature has been focused on the amplitude (i.e.power) of neurofluid fluctuations (Picchioni et al., 2022; Z. Yang et al., 2024), so it remains unclear what the frequency of these fluctuations mean. As mentioned earlier, due to the vasogenic drivers, the infraslow bands (IS1 and IS2) are most strongly associated with ANS control (**Figure 3**). What are the benefits of slow flow fluctuations? While the fast, cardiac-driven pulsations are considered the primary engine for the convective bulk flow that powers the glymphatic system, the slower oscillations in CSF, such as those related to ANS modulation, might have a modulatory effect on this convective flow (Kedarasetti et al., 2022). Moreover, slow-wave flow can maximize the exchange between the CSF and interstitial fluid that enables waste removal. Our work uncovered faster neurofluid fluctuations at hypo- and hypercapnia that are driven at the ANS level (instead of at the arterial-CSF interface), and the implications of this frequency modulation remains to be clarified in future research.

### Limitations

One of the major limitations of this study may be the limited number of participants. However, although our dataset provides unique insight into the physiological activity behind different capnias, only 13 participants were included, which undermines the generalizability of the results from linear mixed effect models. Further, we used a relatively low spatial resolution in order to maximize the sampling rate. While we believe that a high sampling rate is essential for an analysis of this nature, we realize the potential for partial-volume effects in our analysis. As a result of the same reason, aqueduct and fourth ventricle analysis may not be available to all participants due to a limited field of view (also see *method* section). The repetition time (TR=380 ms) we used, despite our efforts to maximize the sampling rate, still poses a challenge to capture all cardiac harmonics.

In our study, we chose ROIs that are distant from the local specific neural activity, such as macrovascular and CSF ROIs. In spite of this, the impact of the capnic on global activity cannot be entirely ignored (F. Xu et al., 2011). There is a possibility that variations in global neural activity and HRV can be transmitted into macrovasculature, which supplies and drains brain blood. In our linear-mixed effect model, these factors have not been included. In addition, some other factors, such as vasomotion and vascular tone, may be difficult to monitor and have not been incorporated into the linear mixed effect model. It is also noteworthy that the macrovascular ROIs used in this study are based on regions with high BOLD power, which may differ from real macrovascular structures based on recent biophysical models (Zhong et al., 2024).

The physiological recording provides insight into cardiac and respiratory activity, and we used them as a surrogate for the activity of the autonomic nervous system. Numerous studies have demonstrated the link between cardiac activity and respiratory activity and ANS activity, however, the modulation of such activities depends on far more than ANS activity alone (Gordan et al., 2015), and such indirect relationships may not exactly replicate ANS activation. On the basis of factors such as healthy young participants and the short capnic period, the ANS appears to be the most likely pathway for cardiac and respiratory variation.

## Conclusions

The study of neurofluid dynamics using in-vivo imaging is an exciting new avenue of research that provides important insights into waste clearance and neuronal signalling in the brain. In this work, we investigate the relationship between capnic condition and neurofluid dynamics as measured by BOLD rs-fMRI signal fluctuations. Inducing different levels of basal CO_2_ has well-established effects on cerebrovasculature, which are in turn linked to the flow of CSF. However, in this study, we found this link is tightly associated with autonomic activity. Moreover, we found that neurofluid dynamics are very similar to those of the global mean signal. We believe these findings can provide new insights into the regulation of neurofluid flow, especially of CSF flow.

## Acknowledgments

The authors would like to acknowledge financial support from Canadian Institutes of Health Research and the Canada Research Chairs Program (JJC) and funding support from Ydessa Hendeles Graduate Scholarship (XZZ).

