## Supplemental Figures for "Modulation of neurofluid fluctuation frequency by baseline carbon dioxide in awake humans: the role of the autonomic nervous system"

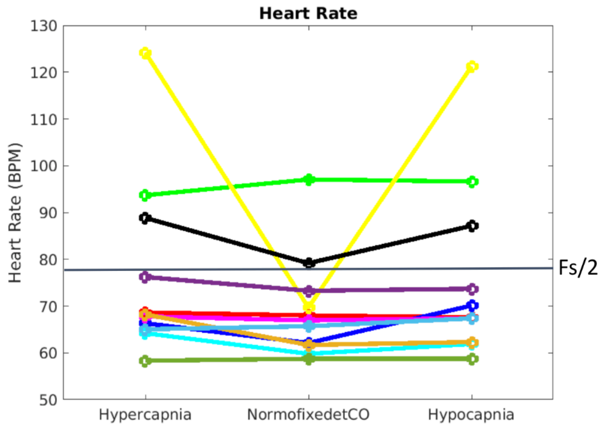

**Figure S1. The participant-level trends across all capnic conditions for HR (left) and RR (Right).** Different colours represent different participants.

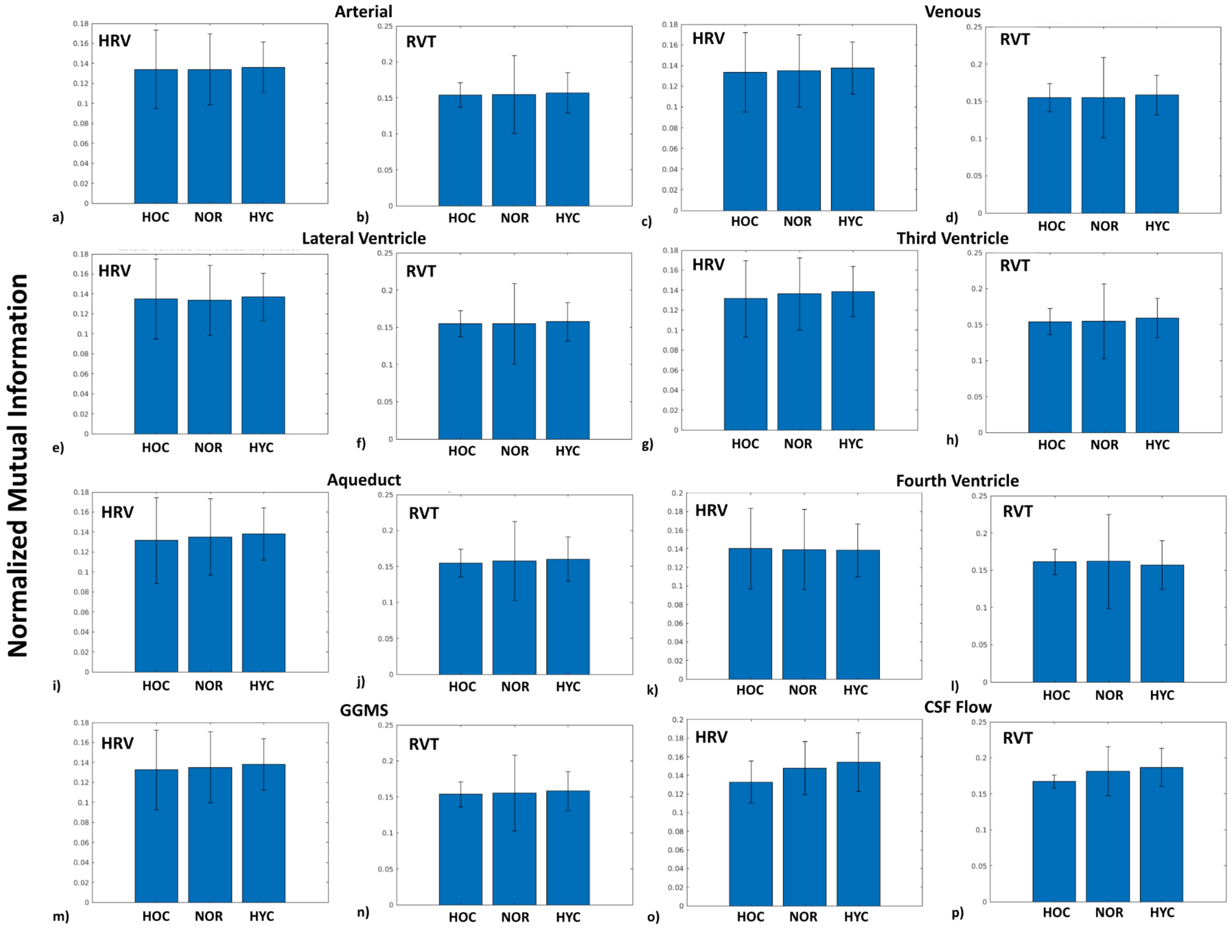

**Figure S2. Mutual Information between HRV, RVT and the ROI-specific BOLD signals across all capnic conditions.** The error bars represent inter-participant standard deviations.

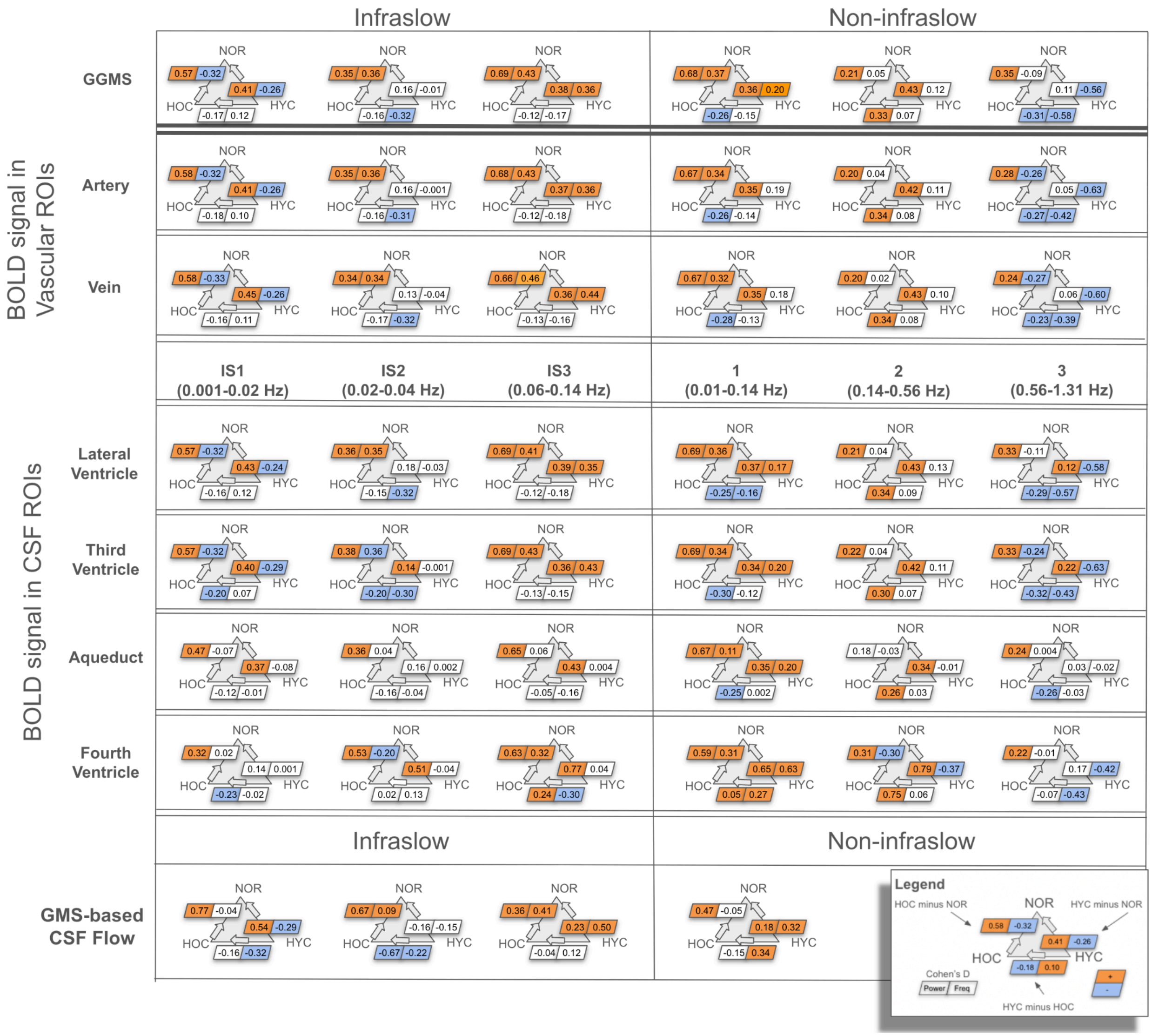

**Figure S3. Effect sizes of differences in rs-fMRI oscillation power and frequency across different capnias.** NOR: normocapnia; HOC: hypocapnia; HYC: hypercapnia. The figure depicts three infraslow bands (IS1, IS2 and IS3) and three non-infraslow bands (1, 2 and 3) for vascular ROIs (artery and vein), CSF ROIs (lateral ventricle, third ventricle, aqueduct and fourth ventricle) and GMS based CSF flow. We have thresholded the D values in order to present only effects greater than 'small'.

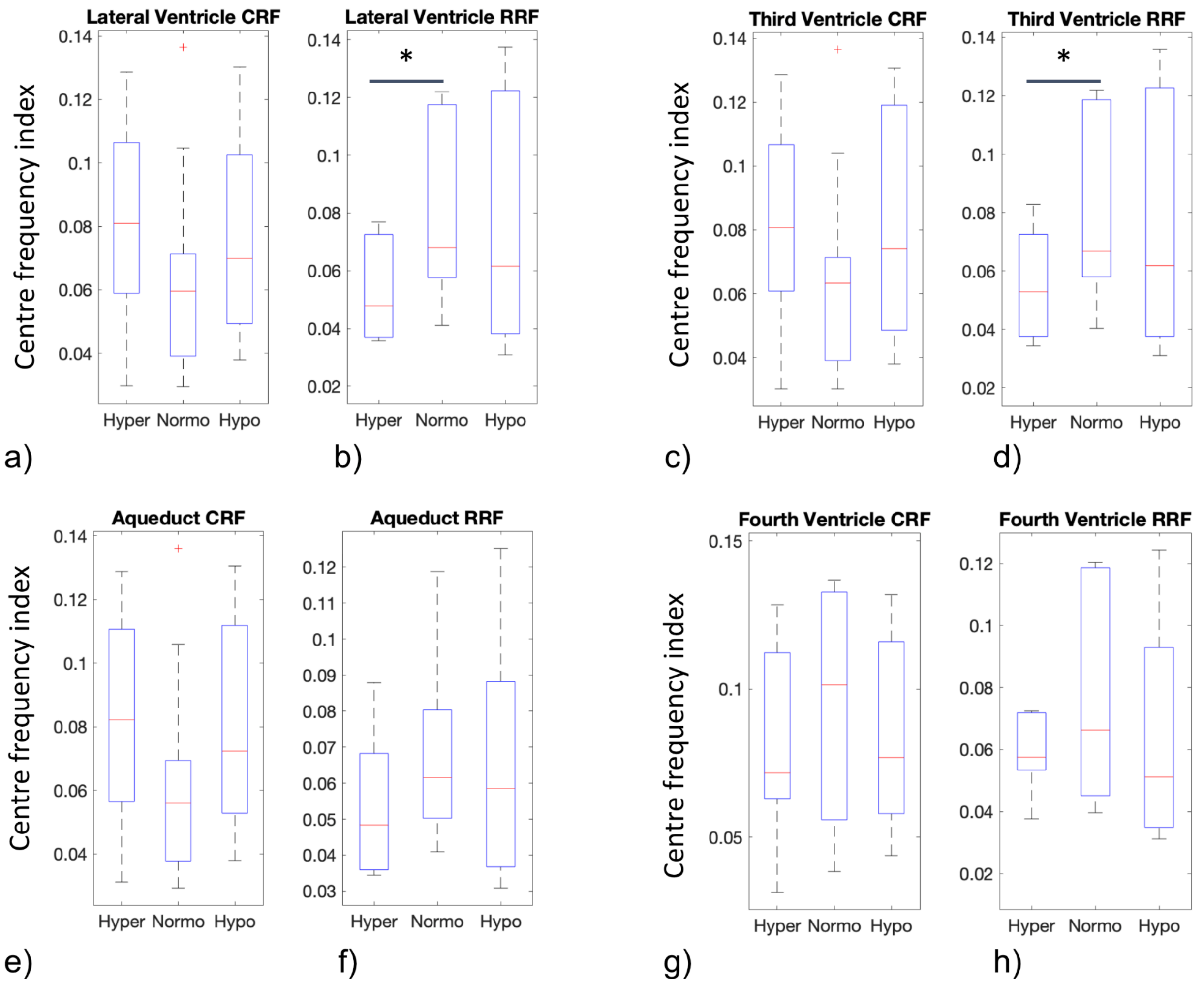

**Figure S4.** **Summarize the centre-frequency indices of RRF and CRF calculated for the rs-fMRI signals in the CSF ROI.**

**Table S1. An overview of rs-fMRI oscillations across different capnias.** Results from neurofluid ROIs and CSF flow are summarized in reference to those from the grey-matter-specific GMS (GGMS). The results were color-coded according to whether BOLD signal power is higher (pink), lower (powder blue), and whether BOLD signal frequency is higher (orange) or lower (blue) in each comparison.

| | Hypocapnia, as compared to normocapnia (HOC minus NOR) | Hypercapnia, as compared to normocapnia (HYC minus NOR) | Hypercapnia, as compared to hypocapnia (HYC minus HOC) |
|---|---|---|---|
| GGMS BOLD | Higher BOLD signal power in all frequency bands<br>Lower BOLD signal frequency in Band IS1 and 3<br>Higher BOLD signal frequency in Band IS2, IS3 and 1 | Higher BOLD signal power in all frequency bands except for Band 3<br>Lower BOLD signal frequency in Band IS1 and 3<br>Higher BOLD signal frequency in Band IS3 | Higher BOLD signal power in Band 2<br>Lower BOLD signal power in Band IS2, 1 and 3<br>Lower BOLD signal frequency in Band IS2 and 3<br>Higher BOLD signal frequency in Band 2 (artey only) |
| BOLD in vascular ROIs | Higher BOLD signal power in all frequency bands<br>Lower BOLD signal frequency in Band IS1 and 3<br>Higher BOLD signal frequency in Band IS2, IS3 and 1 | Higher BOLD signal power in all frequency bands except for Band 3<br>Lower BOLD signal frequency in Band IS1 and 3<br>Higher BOLD signal frequency in Band IS3 | Higher BOLD signal power in Band 2<br>Lower BOLD signal power in Band IS2, 1 and 3<br>Lower BOLD signal frequency in Band IS2 and 3<br>Higher BOLD signal frequency in Band 2 (artey only) |
| BOLD in CSF ROIs | Higher BOLD signal power in all frequency bands<br>Lower BOLD signal frequency in Band IS1 and IS2 (3rd and 4th ventricle)<br>Higher BOLD signal frequency in Band IS2 (lateral ventricle), IS3 and 1 | Higher BOLD signal power in all frequency bands (except for Band 3 in the aqueduct and 4th ventricle)<br>Lower BOLD signal frequency in Band IS1 and 3<br>Higher BOLD signal frequency in Band IS3 and 1 (except for aqueduct and 4th ventricle) | Higher BOLD signal power in Band 2<br>Lower BOLD signal power in all frequency bands except for Band IS3 and 2<br>Lower BOLD signal frequency in Band 1 and 3<br>Higher BOLD signal frequency in Band IS2 (3rd ventricle only) |

| | | | |
|---|---|---|---|
| **GMS -based CSF Flow** | Higher CSF-velocity power in all frequency bands<br>Higher CSF-velocity fluctuation frequency in Band IS2 and IS3 | Higher CSF-velocity fluctuation power in all frequency bands except for Band IS2<br>Lower CSF-velocity fluctuation frequency in Band IS1<br>Higher CSF-velocity fluctuation frequency in Band IS2 and 1 | Lower BOLD signal power in Band IS2<br>Higher CSF-velocity fluctuation frequency in all bands except for Band IS3 |
